# Mechanisms for Communicating in a Marmoset ‘Cocktail Party’

**DOI:** 10.1101/2020.12.08.416693

**Authors:** Vladimir Jovanovic, Cory T Miller

**Affiliations:** Cortical Systems and Behavior Laboratory, Neurosciences Graduate Program University of California, San Diego

**Author notes:** Direct all materials and correspondence to Cory Miller |.

## Abstract

A fundamental challenge for audition is parsing the voice of a single speaker amid a cacophony of other voices known as the Cocktail Party Problem (CPP). Despite its prevalence, relatively little remains known about how our simian cousins solve the CPP for active, natural communication. Here we employed an innovative, multi-speaker paradigm comprising five computer-generated Virtual Monkeys (VM) whose respective vocal behavior could be systematically varied to construct marmoset cocktail parties and tested the impact of specific acoustic scene manipulations on vocal behavior. Results indicate that marmosets not only employ auditory mechanisms – including attention – for speaker stream segregation, but also selectively change their own vocal behavior in response to the dynamics of the acoustic scene to overcome the challenges of the CPP. These findings suggest notable parallels between human and nonhuman primate audition and highlight the active role that speakers play to optimize communicative efficacy in complex real-world acoustic scenes.

## Introduction

Our ability to effectively communicate with others is often complicated by the co-occurrence of other speakers in an acoustic scene, classically illustrated by the Cocktail Party Problem [CPP] ^1, 2, 3, 4^. Studies suggest that humans are able to resolve the challenges of listening in multi-talker scenes using a handful of perceptual cues, including the spatial separation of the speakers and the acoustic idiosyncrasies of individual voices ^4, 5^. Even relatively small distances between talkers can increase intelligibility significantly while differences in each speaker’s voice pitch provides a reliable cue ^6, 7, 8^. In more dynamic scenes involving numerous talkers, these cues may become less clear, requiring listeners to employ top-down perceptual mechanisms to selectively attend to a particular individual’s voice ^9, 10^. During speech, one could learn a talker’s voice and segregate it into a single stream, potentially as a learned schema ^4, 8, 11, 12, 13, 14^, facilitating its segregation from other sounds in the acoustic landscape. Although human and nonhuman primates share the core architecture of the cortical auditory system that distinguish our Order from other taxa ^15, 16, 17, 18^ and face similar challenges of communicating in noisy environments comprising multiple conspecifics vocalizing in concurrence ^19^, there are a notable dearth of experiments testing whether nonhuman primates solve the CPP employing similar mechanisms as humans. In fact, only a handful of experiments have explored whether more general auditory scene analysis mechanisms are evident in our simian cousins ^20, 21, 22, 23^. Certainly observations indicate that primates are able to communicate in noisy environments, but whether this is accomplished because nonhuman primates are talented acoustic scene analyzers that rely primarily on bottom-up auditory mechanism, as is the case in some other nonhuman animals ^3^, or are able to employ more top-down attentional processes characteristic of humans ^9, 24, 25, 26^ is not yet known. Here we implemented an innovative, multi-speaker, interactive playback paradigm that simulates a natural cocktail party while providing experimental control to systematically manipulate multiple dimensions of the acoustic scene to test whether a species of nonhuman primate – common marmosets (*Callithrix jacchus*) – resolves the CPP by employing similar mechanisms as humans. Our goal was not to measure psychoacoustic thresholds of the perceptual processes that reflect auditory scene analysis mechanisms applied generally in audition ^8^, but rather to test how these mechanisms - and potentially others - are leveraged under real-world conditions to overcome the CPP for active communication in a nonhuman primate.

Common marmosets are highly voluble New World monkey who naturally engage in long-distance conversational exchanges within noisy, dynamic communication networks that reflect the CPP ^27^. Like human conversations, the temporal dynamics of these marmoset conversations are governed by learned social rules ^28, 29, 30^. Moreover, marmoset phee calls are individually distinctive and recognizable in conversations ^28, 31, 32^. Building on our previous interactive playback paradigm ^31, 33^, we used a multi-speaker design to construct cocktail parties in which a single live monkey heard phee calls – the species-typical long-distance contact call ^29, 32^ – produced by five Virtual Monkeys (VMs) whose respective vocal behavior differed relative to the subjects. In this innovative design, the behavior of one VM – the Target – was designed to directly interact with the live marmoset, emitting vocalizations in response to the subject in order to engage them in conversational exchanges, while the timing of the other VMs – the Distractors – were independent of the subject. Phee calls from pairs of VM Distractors were constructed in temporal sequences that simulated natural conversational exchanges, such that concurrent conversations between VM Distractor pairs were broadcast in each cocktail party. This innovative paradigm afforded a powerful opportunity to systematically manipulate features of the acoustic scene (e.g. spatial separability and predictability of VM location, distractor density, and the acoustic structure of the vocalizations themselves) in order to explicitly test their effect on subjects’ propensity to engage in conversational exchanges with the Target VM; thus providing key insights into the mechanisms that this primate employs to overcome the challenges of communicating in a cocktail party.

## Results

We tested adult common marmoset monkeys in a series of experiments designed to test how nonhuman primates solve the CPP for active communication. We observed no statistically significant difference in the number of vocalizations produced by subjects across all Test conditions (1-way ANOVA, n = 288, F(15,272)=0.91, p=0.557) suggesting that that all constructed cocktail parties elicited comparable volubility and no overt suppression of vocal behavior in marmosets.

### Baseline Vocal Behaviors

Marmosets perceive calls produced within 10s of their own as a ‘response’ during antiphonal conversations ^28, 33^. Given this generous response window, a critical issue for these experiments was ascertaining whether calls emitted by subjects following a VM call were deliberate responses or simply covaried with the timing of the VM call (i.e. false positive).

To establish false positive rates, we compared marmoset vocal behavior in three conditions – Fixed-Location, Target Baseline and All Baseline. See Methods for a more detailed description of these conditions. Briefly, the Fixed-Location condition involved broadcasting conversations between 2 pairs of Distractor VMs (4 total VMs) and a single, interactive Target VM ^29, 31, 33^. The ‘Target Baseline’ test condition was identical to the ‘Fixed-Location’ condition except that the Target VM calls were not audible. In other words, the timing of a stimulus was recorded but no stimulus was broadcast. This allowed us to ascertain the probability that marmosets’ intrinsic call rate and timing would occur in the response window of the ‘silent’ Target VM calls (i.e. false positive rate) in an environment comprising multiple potential conspecific conversational partners. In the ‘All Baseline’ condition, no vocalization stimuli were broadcast. Similarly to the Target Baseline condition, however, the timing of when interactive Target VM calls would occur was recorded to establish the probability of false positive responses when no calls from other marmosets were broadcast.

We first compared marmoset vocal behavior across these conditions by analyzing subjects’ inter-call interval (ICI), including both conversational exchanges and spontaneous calls. The estimated PDF (see Methods) for the duration of these ICIs was notably different across the three conditions suggesting that the presence of multiple conspecific calls, and their respective behavior affected subjects’ vocal behavior (Figure 2a). More detailed analyses showed that subjects take significantly longer to produce half of all their calls per session (dashed Median data line) in the Fixed-Location condition than the other two conditions (Figure 2b). In fact, subjects’ rate of calling in this condition was relatively constant, while subjects had a bias to produce a higher percentage of calls in the first half of the session for the two baseline conditions – All Baseline and Target Baseline. This was most prominent in the All-Baseline condition which had a significantly different cumulative distribution than the other conditions (n = 18 for each, 95% Confidence Interval).

We next focused on ICI during conversations (i.e. exchanges comprising 2 or more reciprocal call exchanges between the Target VM and subject). Although ‘conversations’ were evident in all contexts (Figure 2c), analyses of the timing of these conversations revealed crucial differences across the test conditions (Figure 2d). Notably, the majority of ‘conversations’ in the All-Baseline condition occurred early each test session suggesting that in the absence of conspecific calls, marmosets were producing phee calls at a high rate during this period, consistent with overall ICI analyses (Figure 2b). Furthermore, while no difference was evident between the Target Baseline and Fixed-Location conditions for approximately the first six minutes of a test session, the occurrence of conversations significantly diverged at this point. Specifically, subjects’ vocal behavior becomes more selective in the Fixed-Location condition.

To further test whether marmosets exhibited a meaningful shift in behavioral strategies after the first six minutes of a test session, we analyzed marmoset vocal behavior in the Target Baseline and Fixed-Location conditions before (Figure 2e) and after (Figure 2f) six minutes of each test session (i.e. >360s). While there was no difference in the PDF of conversation length between these conditions in the first six minutes of each session distributions (Kruskall-Wallis, X^2^(1,n=558)<1e-4, p=0.994; Figure 2E), marmoset conversational behavior significantly diverged after this point (Kruskall-Wallis, X^2^(1,n=990)=17.5, p < 0.0001; Figure 2f). At this time point, subjects’ vocal behavior in conversations are selectively coordinated with the Target VM.This suggests that the first six minutes of a test session are needed to learn the identity of the Target VM, at which point their vocal behavior changes from one of exploration in which they are assessing whether any VMs in the acoustic scene are explicitly interacting with them to a behavioral strategy that is selectively focused on interacting with the Target VM. Based on these analyses, all subsequent comparisons of subjects’ vocal behavior were performed only after the first six minutes of each test session.

### Communication Index

We focused subsequent analyses on marmoset conversations because they reflect a coordinated, reciprocal communication exchange that abides social rules ^30, 34, 35^. Given the high incident of false positive responses in baseline conditions, we developed a single behavioral metric to compare marmoset conversations across the test conditions – the Communication Index (Figure 3a; see also Methods for a more thorough explanation).

Figure 3b plots the results of applying this behavioral metric to marmoset vocal behavior in the Target Baseline and Fixed Location test conditions. There was a main effect on condition and VM as well as an interactive effect (2-Way ANOVA, n=27985, VM: F(4,27975)=572, p<0.0001; Condition: F(1,27975)=54.4, p<0.0001; VM*Condition F(4,27975)=65.3, p<0.0001). Analyses indicated that the Communication Index was significantly higher for the Target VM relative to the distractors in both test conditions (p < 0.0001). As expected, subjects Communication Index for the Target VM was significantly higher in the Fixed-Location condition than the Baseline Target condition (p < 0.0001). Lastly, we applied a Linear Model analysis to the determine whether the Communication Index was modulated by elements of the auditory scene that would indicate its causal relationship with effective communication. This model revealed a significant relationship between Communication Index and the Coefficient of Variation in the Distractor VM ICI (B=-624.79, *t*(111)=-3.83, p=0.000212; Figure 3c, see Methods). In other words, marmosets were able to engage in longer conversations as a function of the predictability of the Distractor VM vocal behavior further indicating that this behavioral metric effectively encapsulates marmosets’ propensity to engage in conversations under these conditions.

### Experiment 1

Here we sought to test how two dimensions of the cocktail party – Distractor Density and Spatial Configuration – affected marmosets’ conversational exchanges. These experiments broadcast 2-Pulse phee call stimuli from each VM at two Distractor Density levels – High and Low – in three spatial configurations – Fixed-Location, Random-Location, and Single-Location (Figure 4a).

Subjects exhibited a significantly higher Communication Index across all conditions to the Target VM relative to Distractor VMs at both the High Distractor (2-way ANOVA, n=9535, VM F(4,9520)=219, p<0.0001; VM*Spatial F(8,9520)=7.11, p < 0.0001) and Low Distractor levels (2-way ANOVA, n=11270, VM F(4,11255)=72.7, p<0.0001; VM*Condition F(8,11255)=4.68, p < 0.0001). Furthermore, at both Distractor Densities, marmosets exhibited a decrease in Communication Index in the Random-Location condition, while showing similar behavior in the other two conditions. The difference in Communication Index with the Random-Location was more modest at the lower distractor density level, as it only reached statistical significance compared to Single-Location, but not in Fixed-Location condition (Figure 4b; p < 0.0001 and p = 0.0726, respectively). At the higher distractor density, however, Communication Index was significantly higher in both the Fixed-Location and Single-Location conditions relative to the Random-Location (Figure 4c; p < 0.0001 and p < 0.0001, respectively). The pattern of results suggests that the predictability of the Target VM in space, rather than spatial separability of the VMs was a key perceptual cue under these conditions.

We next performed analyses to determine whether subjects’ vocal behavior adapted in response to changes in the acoustic scene. Subjects produced a lower ratio of 1 pulse calls at the High Distractor Density level 71.3% to 62.6%, while 2 and 3 pulse calls modestly increased (Figure 4d); a pattern found to be statistically significant (Kruskall-Wallis, X^2^(1,n=4140)=32.7, p<0.0001). Figure 4e further shows that there was a significant change in both the average duration of phee calls (+12.0%), the 1 pulse phee calls (+10.6%), the 3+ pulse calls (+23.3%), but not 2 pulse phee calls, from Low to High Distractor Density (2-Way ANOVA n=4140, Acoustic F(1,4134)=24.9, p < 0.0001; Acoustic*Pulses F(2,4134)=2.39, p < 0.0001). Finally, the median latency within conversations modestly - but significantly - declined from 4.30 to 3.97 (−325 msec) between the Low to High Distractor Densities, respectively (Figure 4f; Kruskal-Wallis test X^2^(1,n=1759)=4.06, p=0.044). These results indicate that subjects increased the median duration of their phee calls while simultaneously decreased their response latency to Target VM calls when communicating at the higher Distractor Density.

### Experiment 2

Primate long-distance contact calls – including marmoset phee calls ^32^ - typically comprise multiple, repeated acoustic pulses to maximize signaling efficacy in noisy environments ^19, 36, 37^. We hypothesized that if redundancy in call structure was perceptually advantageous to marmosets, reducing this characteristic of the call would increase the difficulty of maintaining conversations in some Cocktail Parties. Here we tested subjects in the same Environments as in Experiment 1 but broadcast 1-pulse phee calls from VMs rather than the 2-pulse phee calls used in the previous experiment (see Methods: Test Conditions). Given that subjects already struggled to communicate in the Random-Location condition in Experiment 1, we did not repeat this test condition here.

Similar to Experiment 1, subjects exhibited a significantly higher Communication Index across all conditions to the Target VM relative to Distractor VMs at both the High Distractor (2-way ANOVA, n=7680, VM F(4,7670)=168, p<0.0001; VM*Spatial F(4,7670)=7.98, p < 0.0001) and Low Distractor levels (2-way ANOVA, n=7320, VM F(4,7310)=294, p<0.0001; VM*Condition F(4,7310)=31.2, p < 0.0001). Marmosets exhibited a significantly lower Conversation Index in the Single-Location relative to the Fixed-Location condition at both the Low (Figure 5a; p < 0.0001) and High (Figure 5b; p < 0.0001) Distractor Densities. These results suggest that the spatial separation between the various VMs in the Fixed-Location condition may have afforded perceptual advantages when only hearing 1-pulse phees emitted by the VMs even at the Low Distractor Density level.

Similar to Experiment 1 marmosets’ vocal behavior was affected by the acoustic scene, though the pattern of changes was notably different from the previous experiment. Figure 5c shows that there was a significant change in the distribution of the number of pulses per call made by the subject (Kruskall-Wallis, X^2^(1,n=3030)=32.7, p=0.001). Here, we observed a higher ratio of 1 pulse calls produced by subjects from Low to High Distractor Density (72.1% to 77.5%). These changes did not result in a significant overall change in the duration of calls produced by subjects from Low to High (Figure 5d; 1-Way ANOVA n=3030, F(1,3028)=0.04, p=0.851, overall −0.27%); the significant changes in duration was apparent only when broken down by the number of pulses in the calls subjects produced (2-Way ANOVA Acoustic*Pulses F(2,3024)=7.94, p=0.0004). The 1-pulse calls increased in duration from lower to higher by 5.23% (p = 0.0092), while the 2 pulse and 3+ pulse calls did not change significantly from lower to higher at −1.40% and −13.2%, respectively (p = 0.961 and p = 0.0434). We also observed a significant decrease in latency to respond to the Target VM in conversations at the High Distractor Density level relative to the lower level (Kruskall-Wallis test, X^2^(1,n=1447)=9.5, p=0.0021), similar to Experiment 1, though the latency difference was longer in these conditions (520 msec faster response latency in the High Distractor Density environment, Figure 5e).

### Emergent Acoustic Scene Dynamics Reveal Adaptive Changes in Vocal Behavior

Figure 6a shows the distribution of the mean inter-call interval (ICI) against the calculated High and Low Distractor Density for each session within Fixed-Location and Single-Location. Significant negative correlations exist between the two values for both Experiments 1 and 2 (Pearson’s Linear Correlation: rho = −0.797 & p < 0.0001, rho = −0.928 & p < 0.0001, respectively). The acoustic scene structure revealed by these quantifications emerged because the shorter duration 1-pulse phee calls necessitated a shorter ICI between VM distractor pairs to ensure similar levels of Distractor Density between the experiments. While there are other linear correlations that can be shown, the most significant terms in predicting various behavioral outcomes in our models included the interactive effect of Distractor ICI and Experimental type. Thus, this characterization formed the foundation for the subsequent statistical analyses aimed at characterizing the relationship between the emergent scene structure and marmoset vocal behavior in these experiments.

We applied a linear model to test how facets of marmoset vocal behavior covaried with dimensions of the acoustic scene. The following were input into the Linear Model – VM Pulse # (2-pulse:Expt 1, 1-pulse:Expt2), Low and High Distractor Density, and Fixed and Single conditions – for a total of 144 sessions. We also chose to include the calculated Distractor Density for each session along with the Distractor ICI (see Methods). Given a strong positive correlation between Distractor ICI and standard deviation (Pearson’s Linear Correlation: rho = 0.931 and p < 0.0001), we took the coefficient of variance (COV, standard deviation divided by mean) as a way to encapsulate these two correlated factors while avoiding rank deficiency in any linear model (COV v Mean ICI, Pearson’s Linear Correlation rho = −0.0980, p = 0.243. Figure 6b). This also gave an added benefit of enumerating the relative dispersion of the Distractor ICI. This analysis yielded six total predictor variables.

We tested eight interactive linear models which included 22 terms (1 intercept, 6 linear predictor terms, and 15 pairs of distinct predictor terms). The statistical threshold for significant terms and models was corrected for multiple comparisons with Bonferroni correction based on 22*8=176 comparisons with a corrected P value threshold at 0.05/176 = 0.000284. Of these eight models, three models reached significance: Calls Produced (F(21,111) = 3.88 with adjusted R^2^=0.314), Conversation Count (F(21,111)= 4.19 with adjusted R^2^=0.337), and Communication Index (F(21,111) = 5.47 with adjusted R^2^=0.415). One significant term was shared across the three models: The Distractor ICI x 1/2 Pulse (Experiment). COV Distractor ICI x Distractor ICI (which results in standard deviation Distractor ICI) was significant only for Communication Index. Figure 6c-f plots the four significant terms against the respective response variables in interaction effects plots. Each image plots the adjusted response function of the given response variables on the Y-axis against the values of the first predictor in the interactive term with the second predictor at fixed values (for categorical: all levels, and numeric: minimum, maximum, and average of minimum and maximum). Given that all four interactive terms have significant coefficients within their respective models, and that the slopes of the lines in all four plots are not parallel, there is significant interactive effect between the predictors for predicting the number of number of calls produced by the subject, the conversations made in a given session, and the mean Conversation Index with respect to Target VM and subject.

Presenting subjects with VM calls comprising either 2 or 1 pulse phee calls – Experiments 1 and 2, respectively – resulted in opposite effects on the adjusted response variables. For Conversation Count (Figure 6c), Calls Produced (Figure 6d) and Communication Index (Figure 6e), these behavioral metrics revealed a positive correlation with Distractor ICI in Experiment 1, but a negative relationship in Experiment 2. In other words, when hearing 2-pulse VM calls in Experiment 1, subjects were more likely to produce more calls, engage in more conversations, and have higher Communication Index values as the Distractor ICI increased in duration. By contrast, the opposite was true when hearing only 1-pulse phee calls in Experiment 2. In other words, different behavioral strategies were needed to optimize communication based on the specific dynamics of the scene.

A further significant factor affecting marmoset vocal behavior in the linear model was COV Distractor ICI (Figure 6f). As the Distractor ICI increased, at low COV, the mean Communication Index decreased. At the highest level of COV, the opposite relationship emerged with increasing Communication Index (with a smaller relative change). This suggests that as the predictability of the Distractor ICI increased (high to low COV); shorter Distractor ICI were optimal for the subject to produce calls and engage in more conversations with the Target VM. Similarly, pertaining to the importance of spatial predictability for marmosets in Experiment 1, temporal predictability was advantageous for marmosets to navigate the complex acoustic scene and selectively engage with the Target VM.

## Discussion

Here we employed an innovative multi-speaker paradigm to construct real-world cocktail parties and test how a New World primate – common marmosets – resolves the challenges of these acoustic scenes for active communication. We report that marmosets not only demonstrated a remarkable ability to overcome the experimental perturbations imposed on them and engage in conversational exchanges but did so by complementing mechanisms of audition – similarly to humans ^2^ – with adaptive modifications of their own vocal behavior. These findings suggest that elucidating the neural mechanisms that underlie the CPP in human and nonhuman primates may also need to consider that listeners are active explorers of the world who actively modify their behavior in response to the changing features of the acoustic scenes to optimize communication rather than rely solely on audition.

Engaging in conversational exchanges in these cocktail parties likely relied on a schema-based learning mechanism for speaker stream segregation ^8, 11, 13^. First, the identity of the Target VM needed to be learned in each session. While the spectro-temporal structure of marmoset phee calls is relatively stereotyped, each monkey’s phee is individually distinctive and perceptually recognizable ^31, 32^. As a result, segregating one caller’s phee call from amongst many conspecific vocalizations presents a distinct challenge that relies on learning the identity of an interactive conversational partner. Second, learning the identity of the Target VM was based on its distinctive vocal behavior. While subjects heard high numbers of calls from Distractor VMs in all conditions, only the Target VM vocal behavior was designed to be interactive with subjects ^28, 29, 33^. Therefore, marmosets learned the identity of the Target based on the statistical occurrence of VM Target calls relative to their own rather than anything intrinsic to the vocalizations themselves. Indeed, evidence suggests that this process took time, as subjects needed ~6mins of a test session to learn the identity of the Target VM (Figure 2d-f). Third, once the Target VM identity was learned, marmosets also needed to continuously monitor that conspecifics’ behavior in order to coordinate their own relative vocal behavior for conversations. Marmoset conversations abide social rules that govern the temporal dynamics of these interactions ^28, 29, 30, 33^, but the periodicity of these exchanges is notably slow. The median interval between conspecific calls in conversations is ~3s, but it can range up to 10s ^29, 33^. The cacophony of marmoset phee calls broadcast in these experiments – particularly at the high Distractor Density level – created a particularly challenging environment in which to perceptually track the Target VM. Evidence suggests that marmosets relied on a reliable spatial cue to focus attention and implement a schema-based mechanism to solve the CPP.

The pattern of results suggest that marmosets employed auditory attention to resolve the CPP. Experiments in humans involving multiple speakers found that when the spatial position of each talker randomly changed across locations, subjects’ intelligibility scores decreased ^7^. Likewise, human subjects performed significantly better when the spatial location of the target was cued prior to hearing the sound ^38^. In both cases, it was concluded that the predictability of a talker’s position in space allowed subjects to focus attention to that position in space. When that predictability was eliminated, attention could not be focused, and it accordingly had a negative impact on subjects’ capacity to understand what was spoken. The Random-Location Condition in Experiment 1 was designed to test whether a similar pattern would emerge in marmosets. Importantly, the vocal behavior of the VMs (i.e. the acoustic scene) was identical across all three spatial conditions, and the only difference in the Random-Location condition was a lack of predictability for the location from which each phee call was broadcast. As shown in Figure 4b, subjects’ exhibited lower Communication Index in Random-Location condition than the other two spatial configuration, an effect that increased when Distractor Density was higher (Figure 4c). These results suggest that the spatial predictability, rather than the spatial separability, of the VM callers was key to resolving the CPP under these conditions suggesting that, like humans ^7, 38^, attentional mechanisms were likely necessary to learn a schema for speaker-stream segregation.

Results from Experiment 2 contrasted with Experiment 1 in several important ways that may reveal an evolutionary relationship between vocal signal design and audition in marmosets. Nonhuman primate long-distance contact calls – including the marmoset phee – often comprise the repetition of a single syllable, a signal design feature conjectured to limit degradation of the signals communicative content when transmitting long distances through noise acoustic environments ^36, 37, 39^. Marmoset phee calls, for example, consist of 1-5 acoustically similar pulses ^32^. While marmosets performed similarly in the Fixed- or Single-Location conditions when hearing 2-pulse phee calls in Experiment 1 (Figure 4b,c), marmosets struggled to engage in conversations when only 1-pulse phee calls were broadcast from a Single-Location in Experiment 2 (Figure 5a,b). In other words, under these conditions spatial-release from masking was necessary to identify the Target VM and maintain conversational exchanges ^40, 41, 42^. By reducing the number of pulses in each call, we effectively halved the amount of acoustic information available to both identify the Target VM and recognize it in the cocktail party for subsequent potential interactions. Indeed, reducing the number of pulses in the contact calls of closely related tamarin monkeys significantly impaired their ability to recognize the caller’s identity ^43^. This suggests that the acoustic redundancy of a two-pulse phee call is crucial to maintaining active conversations in noisy environments because it provides necessary information about the caller’s identity. Selection for multi-pulsed phee calls in marmoset evolution, and potentially more broadly for other nonhuman primates ^36^, may have been driven by the limits of audition for parsing vocalizations and recognizing callers amid the myriad of biotic and abiotic noise common in the species forest habitat.

Results from a Linear Model indicated that these primates did not solely rely on audition to effectively communicate in cocktail parties, but adaptively change their behavior in response to the dynamics of the acoustic scene ^44, 45, 46, 47, 48^. To control for acoustic interference, it was necessary to decrease the inter-call interval (ICI) between phees in the Distractor VM conversations which resulted in a systematic change in the periodicity of the Distractor VM conversation (i.e. variance and inter-call interval). The effect of these cocktail party characteristics was a tactical change in marmoset vocal behavior based on whether they heard 2-pulse (Experiment 1) or 1-pulse (Experiment 2) phee calls. When Distractor VM Conversations comprised 2-pulse phee calls in Experiment 1, marmosets produced more calls (Figure 6c), more conversations (Figure 6d), and resulted in an increased Conversation Index (Figure 6e) when Distractor ICI increased. In stark contrast, marmosets exhibited the opposite effect in Experiment 2 when hearing Distractor VM conversations comprising 1-pulse phee calls, biasing all three facets of vocal behavior to shorter Distractor ICI (Figure 6c-e). In other words, optimizing communicative efficacy relied on a different strategy depending on the call variants produced by the VMs in the particular acoustic scene. A second adaptive behavioral strategy that emerged from this model was the influence of the predictability of Distractor VM call timing. Marmoset overall calling was inversely related with the variance of the Distractor VM ICI (Figure 6f) indicating that they were significantly more likely to increase Conversation Index when they could reliably predict the timing of the Distractor VM calls. These patterns of behavior suggest that marmosets are not simply treating the Distractor VM calls as a broad masker; instead, they are attending to the dynamics of those conversations as well as their own. In other words, marmoset attention appears to be divided between the Target VM and Distractor VM vocal behavior to resolve the challenges of the CPP.

The challenges of communicating in cocktail parties is a daily occurrence for human and nonhuman primates. Results presented here suggest that solving the cocktail party in real-world, multi-talker environments may be a far more dynamic, active process than is typically considered ^7^. Notably, the unique insights reported here were possible because of the innovative multi-speaker paradigm developed for these experiments to construct cocktail parties and systematically manipulate key properties of these social and acoustic landscapes. A broader implication of these findings is the opportunity to leverage this exciting paradigm to investigate the neural basis of these perceptual and cognitive mechanisms underlying the CPP in the primate brain. Neural recordings in human auditory cortex have highlighted the role of attention in representing speakers in complex acoustic scenes comprising multiple talkers ^49, 50^, but relatively little remains known about how other neural substrates in the auditory system contributes to the myriad of related processes. Marmosets share the core functional architecture of the auditory system with all other human and nonhuman primates ^16, 18, 51, 52^, and has been a key primate model of sound processing, including vocalizations, for many years ^27, 53, 54, 55, 56^. By integrating existing technologies for recording neural activity in freely-moving marmosets with the current behavioral paradigm reported here, the potential to explicate the circuit level mechanisms in the primate brain that underlie the CPP can be realized.

## Methods

### Subjects

Ten adult marmosets participated as subjects in this study. Six marmosets (3 females and 3 males) were subjects in Experiment 1 and 2 from September 2019 to May 2020. Two of these subjects (1male and 1 female) as well as four additional adult marmosets (2male and 2 female) served as subjects in the All Baseline condition in March 2021. All marmosets were social housed in pair-bonded family units that comprised of two adults, and up to two generations of offspring. The UCSD Institutional Animal Care and Use Committee approved all experimental procedures.

### Experimental Design

All experiments were performed in a ~4 × 3 m Radio-Frequency Shielded testing room (ETS-Lindgren). Individual subjects were transported from their home cage in clear acrylic transport boxes to the experimental chamber and tested individually. Subjects were placed in an acrylic and plastic mesh test cage (32 × 18 × 46 cm) designed to allow the animals to climb and jump freely along the front wall of the cage similarly to previous experiments ^28, 31^. The cage was placed on a rectangular table against the shorter side of the room. Seven speakers (Polk Audio TSi100, frequency range 40-22,000 Hz) were placed on the opposite side of the room arranged to maximize distance relative to all other speakers in both the horizontal and vertical planes (Figure 1A). All vocal stimuli were broadcast at 80 dbSPL as measured 0.5 m in front of the speaker. A cloth occluder divided the room to prevent the subjects from seeing any of the speakers during testing. One directional microphone (Sennheiser, model ME-66) was placed approximately 0.3 m in front of the subject to record all vocalizations produced during a test session. Another directional microphone was placed in front of the central speaker as well. We tested subjects three times to each test condition across two experiments while randomized. The order of each condition within the individual Experiments was counterbalanced across subjects in a block design for the High and Low Distractor Density levels.

**Figure 1.**
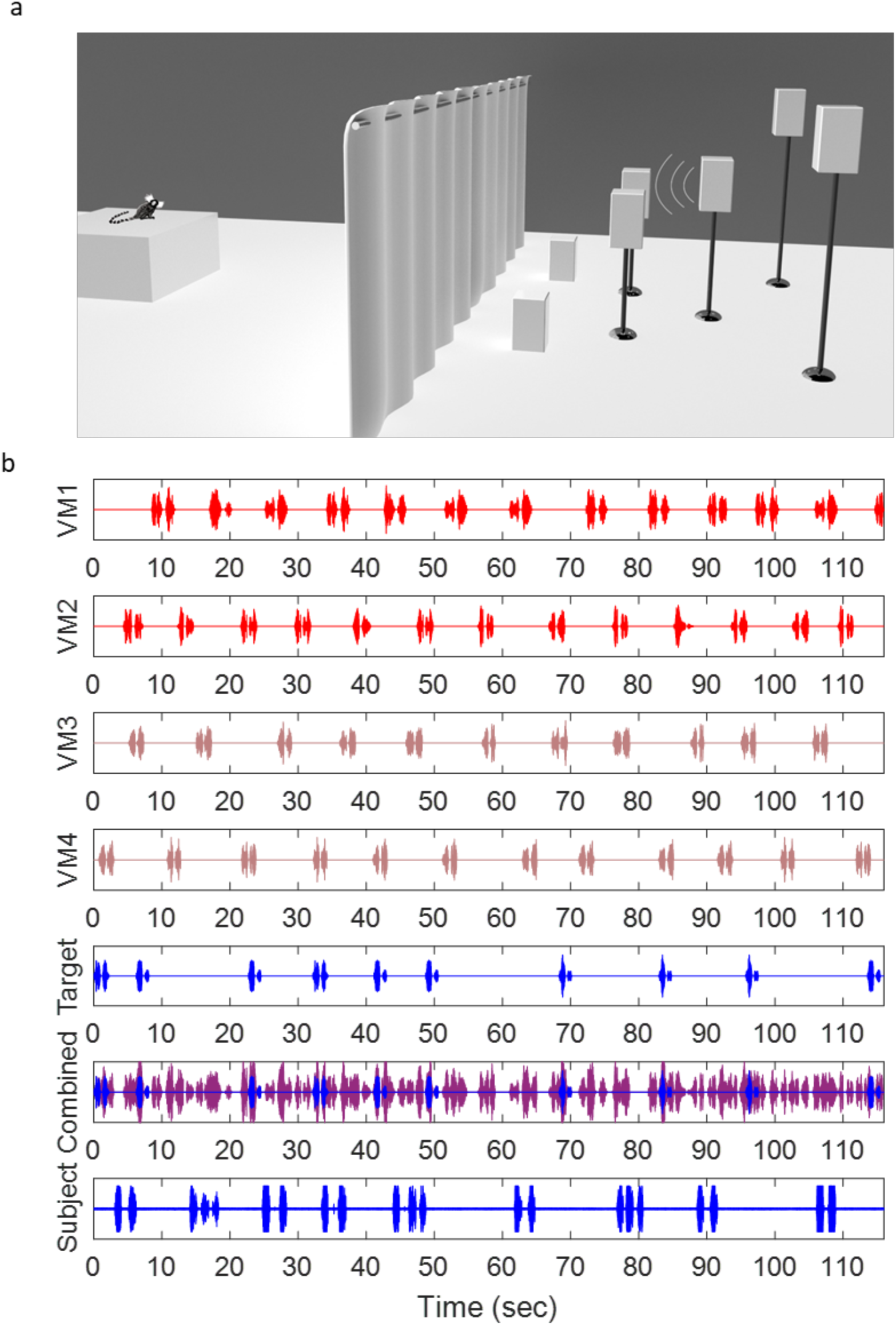
Design of the marmoset Cocktail Party experiments. (a) Schematic drawing of the spatial configuration of the testing room. Subjects were placed in a clear acrylic box with a mesh front (box around subject not pictured). Seven speakers were positioned to have spatial separation in height, distance and width. An opaque curtain was placed equidistant between the subject and speakers to occlude visual access. (b) An exemplar two-minute sample of the vocalizations broadcast by the Virtual Monkeys (VM) and a live marmoset subject from a High Distractor Density, Fixed-Location session in Experiment 1. VM 1-4 are Distractors. VM1 and VM2 (shown in red) have been designed to broadcast 2-pulse phee calls that reflect a conversation with each other, while VM3 and VM4 (shown in brown) are likewise designed to engage in a reciprocal conversational exchange. The Target VM (blue) is engaged with the live marmoset Subject in an interactive reciprocal exchange based on subjects’ vocal behavior. The combined view shows the summation of all VM phee calls – Distractors (purple) and Target (blue).

**Figure 2.**
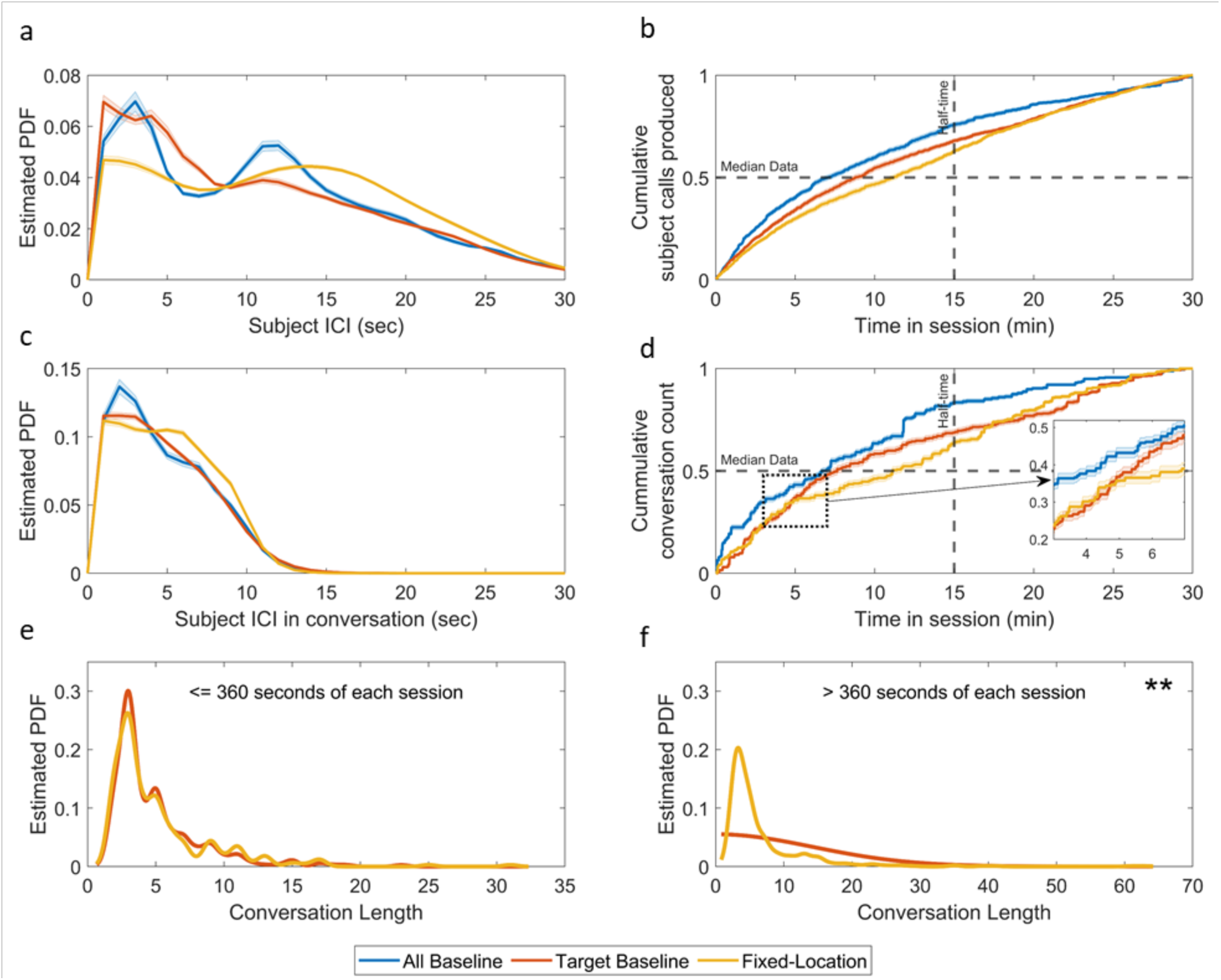
Comparison of marmoset vocal behavior in three ‘Baseline’ conditions: All Baseline, Target Baseline and Fixed-Location (n = 18 for each). (a) Distribution of subject inter-call interval from offset to onset of subsequence subject calls that were spontaneous or the initiations of conversions. 95% CI in shaded areas. (b) Cumulative distribution of subject calls produced normalized for comparison across baselines. 95% CI in shaded areas. Median data refers to 50% of all calls produced by the subject in a session. Half time refers to half of the duration of a session. (c) Distribution of subject inter-call interval only within conversations that contain at least two subject responses. 95% CI in shaded areas. (d) Cumulative distribution of conversations counts as mentioned before. Insert shows an expanded view from to 3 to 7 minutes. 95% CI in shaded areas. (e) Estimated PDF for all conversation lengths of at least 2 or more subject calls for the first six minutes of all sessions. (f) Estimated PDF for all conversation lengths of at least 2 or more subject calls after the first six minutes of all sessions (> 360s). ** p-value < 0.001.

Cocktail parties were constructed using an innovative multi-speaker paradigm in which vocalizations were broadcast from five, software generated Virtual Marmosets (VMs) (Figure 1A). The unique individual identity of each VM was determined by (1) broadcasting prerecorded vocalizations from an individual marmoset in the UCSD colony and (2) its vocal behavior relative to the live subject and other VMs. With respect to this later characteristic, VM vocal behavior was determined by their designation as a Target or Distractor. Similar to our previous experiments ^28, 31^, the behavior of Target VM was specifically designed to directly engage subjects in the species-typical natural conversational exchanges by utilizing an interactive playback design. To this end, the Target VM would broadcast a phee call response within 1-5s with an 85% probability each time subjects produced a phee call. In successive vocal exchanges between the subject and target (e.g. a conversational exchange), the Target VM would broadcast a response with 100% probability to maintain the vocal interaction. If subjects did not produce a call within 15-30s, the Target VM would broadcast a spontaneous call. Custom-designed software recorded vocal signals produced by the test subject from the directional microphone positioned in front of the animal and identified when subjects produced a phee call. By contrast, the timing of Distractor VM phee calls were independent of subjects’ behavior, occurring at a predetermined interval. In each test condition, we generated two pairs of Distractor VMs. Each pair was designed to directly engaged each other in conversational exchanges. The timing of phee calls within these conversations was determined by the parameters of the test condition.

#### VM Stimulus Sets

All phee calls used as stimuli in these experiments were recorded from animals in the UCSD colony using standardized methods in the laboratory described in previous work ^28, 31^. Briefly, two monkeys were placed in separate testing boxes positioned ~3m from each other with an opaque cloth occluder located equidistant between the boxes to eliminate visual contact between the animals. Directional microphones (Sennheiser ME-66) were placed directly in front of each subject to record vocal output separately from each animal. Naturally produced calls were recorded direct to disk over a 30min session. At the conclusion of the session, custom-designed software was used to extract two-pulse phee calls produced during each session. Phee calls produced within 10s of a conspecific phee were classified as ‘antiphonal’ responses, while those produced after this threshold were classified as ‘spontaneous’ phee calls. These designations were based on previous research ^33^. Each VM in a test session would only broadcast antiphonal and spontaneous phee calls from a single marmoset. The stimulus sets used as the basis for all Target and Distractor VMs was randomized across test sessions. The VMs stimulus sets used to construct each cocktail party were never produced by animals in a subject’s home cage because of confounds that might occur due to social relatedness ^29^. Although marmosets naturally produce phee calls comprising 1-5 acoustically similar pulses, the modal call variant is the 2-pulse phee ^32^.

### Test Conditions

We selectively manipulated two dimensions of the scenes - *Spatial Configuration* & *Distractor Density* - to directly test their respective impact on how marmosets resolved the challenges of communicating in a cocktail party in two separate experiments distinguished only by the phee call variant broadcast to subjects. Experiment 1 tested subjects using two-pulse phee calls as vocalization stimuli produced by VM, while Experiment 2 broadcast only 1-pulse phee calls from the VMs. The 1-pulse calls were created by removing the second pulse in the 2-pulse phee call repertoire of all the VMs. In general, this would mean half the duration of a the standard 2-pulse call played in Experiment 1. Subjects were tested three times on each Spatial Configuration at each Distractor Density. The order of the trials was randomized and counterbalanced across subjects.

#### Spatial Configuration

The spatial location of the VMs was manipulated by broadcasting the phee stimuli in three different speaker configurations: *Fixed-Location, Random-Location* and *Single-Location* (Figure 3A). These configurations allowed us to contrast the effects of spatial separation between the callers and the predictability of a caller’s position in space on marmoset vocal behavior.

**Figure 3.**
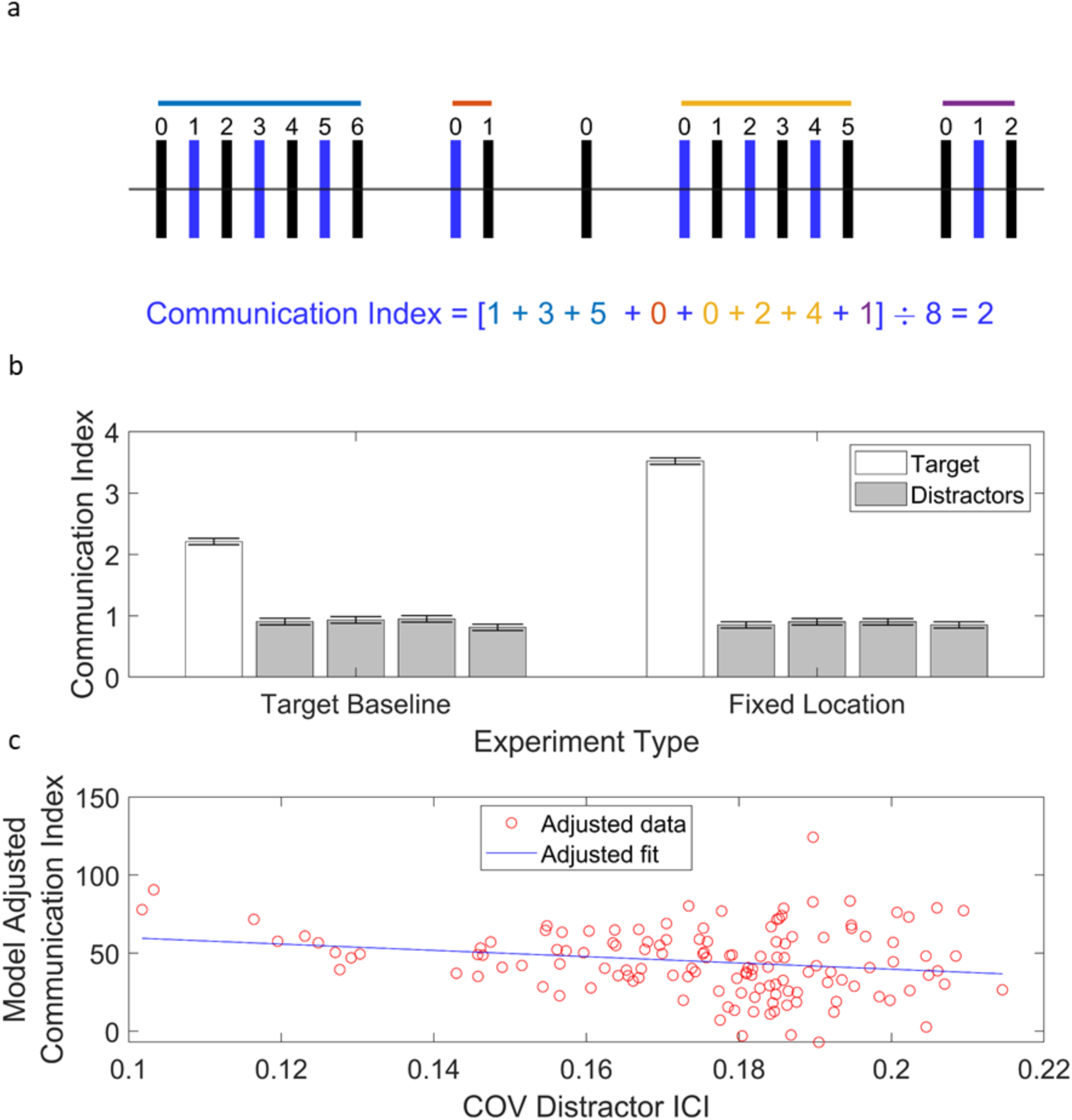
Communication Index. (a) Schematic showing an abstract exchange of phee calls between a VM and a subject. Black bars indicate calls produced by the Target VM while blue bars indicate calls produced by subjects. Colored horizontal lines above indicate vocal exchanges with varying lengths (6, 1, 5, 2). Each call produced by the subject within a vocal exchange is labeled by zero-based numbering. These values are summed and divided by the total number of calls produced in the session. (b) Bar plot showing the calculated Communication Index distributions in comparison to each VM across the Target Baseline and Fixed Location conditions. Error bars represent 95% Confidence Interval and multiple comparison corrected. Target VM differences were significant at p < 0.0001. (c) Linear Model outcome shows a significant relationship between the predictability of Distractor VM calls (Coefficient of Variance of the Distractor Inter-call Interval (ICI)) and Communication Index (B = −624.79, *t*(111)=-3.83, p=0.000212).

**Figure 4.**
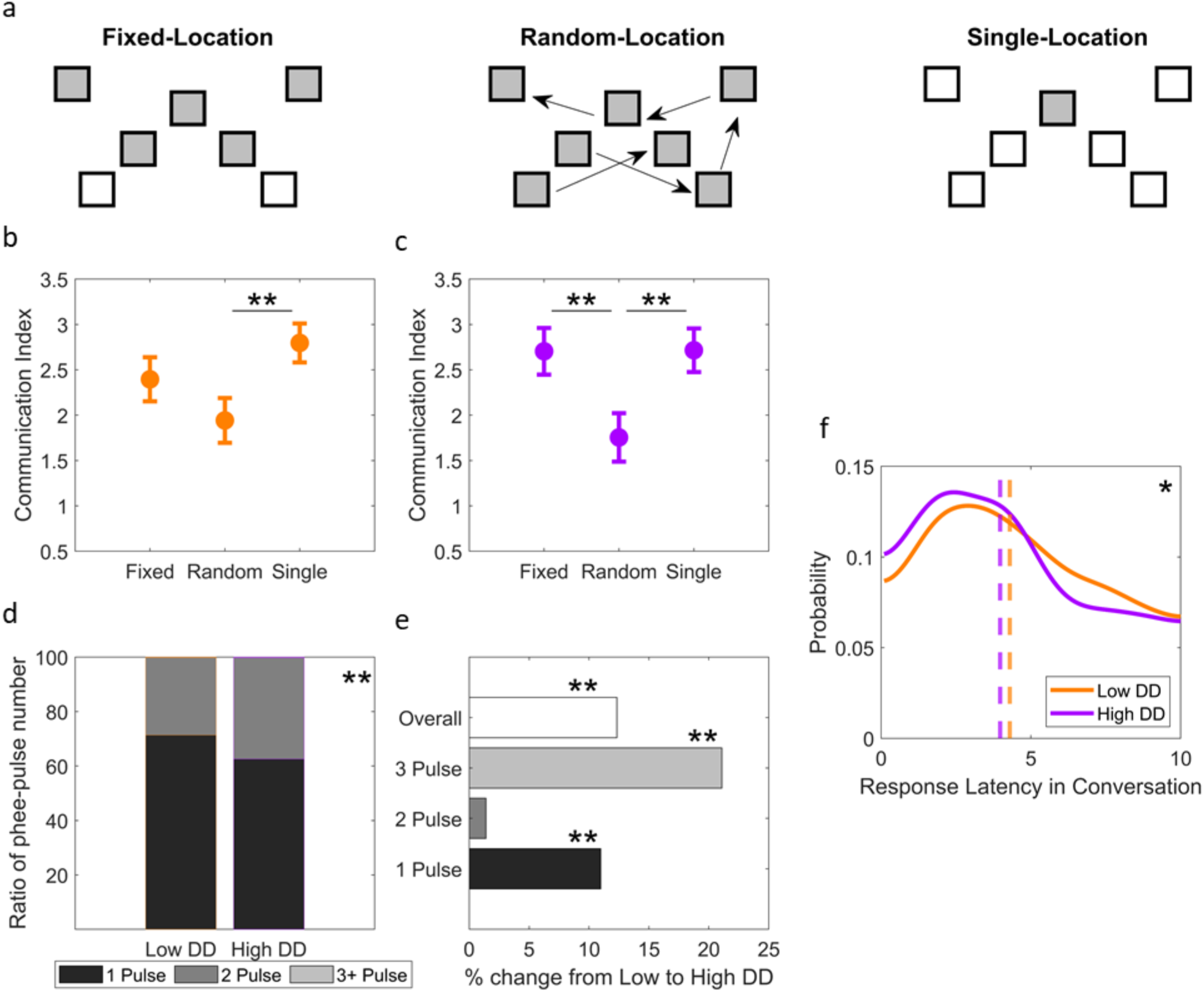
Results from Experiment 1. (a) A schematic drawing of the spatial configuration of the seven speakers used in these three test conditions from above: Fixed-Location, Random-Location, and Single-Location. Grey shading indicates which speakers broadcast phee calls for that condition. Arrows in the Random-Location condition indicate the fact that the speaker location from which each VM phee was broadcast was randomized for each stimulus presentation across the seven-speakers. (b, c) Plots the Mean Communication Index [95% CI] for Fixed-Location, Random-Location, and Single-Location test conditions. ** Significant difference between two conditions, p < 0.0001. (b) Plots Communication Index for the Low Distractor Density condition, while (c) plots results from the High Distractor Density condition. (d) Stacked bar graph showing the distribution phee calls produced by subjects that comprised 1-Pulse (black), 2-Pulses (dark grey) and 3 or more pulses (light grey, though too small to see) in both the Low DD and High DD environments. ** Significant difference between distributions, p < 0.0001 (e) The change in duration of all calls, and sub-groups of phee-pulse calls from Low to High Distractor Density (DD) is shown as percent change. ** Significant difference for that category, p < 0.0001. (f) Estimated PDF of subjects’ latency to respond to the Target VM in conversations in both Low DD (red) and High DD (blue) conditions. The median value is shown as a dashed vertical red bar – Low DD – and blue bar – High DD. * Significant difference between distributions, p < 0.05.

##### Fixed-Location

In this configuration, the calls of each VM were broadcast from among five distinct, spatially separated speakers. This scene afforded subjects spatial separability of each VM from a consistent spatial location for the duration of the experiment. Because this configuration provided the most consistent perceptual cues to subjects, we also used data from this condition for comparison of subjects vocal behavior with the Baseline conditions described below.

##### Random-Location

Like the Fixed-Location condition, VM calls were broadcast from distinct spatially separated speakers. Rather than each VM broadcast from their own speaker for the duration of the experiment, speaker location was randomized across all 7 potential speakers during each broadcast. No VM call would be broadcast from the same speaker twice in a row, nor was there any overlap in VM calls from the same speaker. As a result, subjects were afforded spatial segregation of the VMs, but with no predictability for where the VM would emit a call.

##### Single-Location

Here all VM stimuli were broadcast from a single speaker, thereby eliminating spatial separation of the different callers.

#### Distractor Density

Distractor Density was calculated as the ratio of the Target VM calls that temporally overlapped with Distractor VM calls. This property was manipulated to two levels – Low and High – by changing the relative inter-call interval between phees broadcast between VM Distractor pairs. In the ‘Low’ distractor density scene (~70% acoustic overlap), Distractor VM conversations had an inter-VM call interval ranging 1 to 3.5 sec in Experiment 1 [2-pulse phee calls] and 1 to 2.5 sec in Experiment 2 [1-pulse phee calls]. In the ‘High’ distractor density scenes (~90% acoustic overlap), Distractor VM conversations had an inter-VM call interval ranging from 0.5 to 1.0 sec in Experiment 1 [two-pulse phee calls] and 0.5-0.75 sec in Experiment 1 [one-pulse phee calls]. The shorter inter-VM call interval ranges for Experiment 2 were used to maintain the same level of Distractor Density when the shorter one-pulsed phee calls were used as stimuli.

### Baseline Conditions

Because the long time window over which marmosets perceive calls from conspecifics as a response to their own (10s) ^33^, this condition was designed to test the probability that subjects will emit vocalizations at times consistent with a vocal response to an actual call (i.e. false positive). Subjects’ vocal behavior under these conditions could, therefore, be compared to the Fixed-Location to ascertain the which properties were most characteristic of active communication.

#### Target Baseline

The following condition was performed to establish the probability of false-positive responses when marmosets were in Cocktail Party environments comprising multiple conspecific callers. Subjects were tested in an environment identical to the Fixed-Location condition with one key exception. Rather than broadcast the calls of the interactive Target VM, here those vocalizations stimuli were not audible to subjects. Rather than broadcast the stimulus, the timing of that stimulus was recorded in the event log. This allowed us to quantify marmoset vocal behavior in the same dynamic acoustic scenes as they experienced in the Test Conditions, but without an interactive conversational partner. Subjects were tested three times in the Target Baseline trials for both High and Low Distractor Densities in Experiments 1 and 2. These trials were randomized and counterbalanced with the Test Condition trials.

#### All Baseline

Our initial experiment tested subjects only in the Target Baseline condition. We later determined that quantifying subjects’ vocal behavior in the absence of any conspecific calls would be important to determine how marmoset vocal behavior differed in the Cocktail Party environments relative to when they heard no conspecifics. These trials were identical to the Target Baseline condition except that we did not broadcast the Distractor VM calls. In other words, subjects heard no marmoset calls. We tested six subjects three times on this condition.

#### Statistics

A two-tailed One-sample Kolmogorov-Smirnov test was used to inspect most data for normality like Communication Index. N-way ANOVAs (1,2, and 3) were performed on data sets using the anovan function in MATLAB. If there were significant main or interactive results, post-hoc multiple comparisons were corrected by Tukey’s Honestly Significant Difference Procedure within the multcompare functionin MATLAB. For distribution tests, we used Kruskal-Wallis. 95% confidence intervals were two-tailed [0.025, 0.975] based on standard error. Bresuch-Pagan was used to test heteroskedasticity of residuals. Normality tests of the residuals used the Lilliefors test.

### Data Analysis

We calculated the following behavioral metrics to quantify changes in subject vocal behavior relative to the Target and Distractor VMs as well as standard acoustic parameters, such as call duration and response latency.

#### Communication Index

Our analyses focused on marmoset conversations to explore how these monkeys solved the CPP because this social behavior is indicative of an active, coordinated communication exchange between marmosets ^28, 30^. Previous experiments in marmosets determined that phee calls produced within 10s following a conspecific phee call were perceived as a ‘response’ to the initial call by conspecifics and were significantly more likely to elicit a subsequent vocal response, while those produced after this threshold did not elicit vocal responses from conspecifics ^33^. We defined a conversation as each behavioral epochs in which two individuals engage in a series of alternating, reciprocal phee exchanges during which the inter-call interval between conspecific phee calls is <= 10s ^28, 57^. Each conversation ended when the subject did not produce a phee call within 10s of the offset of the preceding Target VM call.

We calculated a Communication Index to quantify the relationship between phee calls produced in conversations weighted by its length relative to all phees produced by the subject in a session (Figure 3A). By adopting a single behavioral metric, we were able to directly compare subjects’ behavior across different test conditions. To calculate the Communication Index, we first identified all instances of phee calls produced by subjects in a test session. Subjects calls in these conversational exchanges were assigned a number based on their linear order in the vocal exchanges sequence. In other words, the first response was assigned 1, the second successive response was assigned 2, etc. Spontaneously produced calls and phees produced as the initiating call of a conversation by subjects were assigned 0. These numbers were summed and divided by the total number of phee calls produced in the session (Figure 2B).

#### Interference Ratio

We measured the temporal overlap between the Distractor VMs calls and the Target VM calls to determine the amount of acoustic interference that occurred. Each time a Target VM call was broadcast, we measured the duration of time it temporally co-occurred with the duration of any Distractor VM call. The resultant ratio indicates the percentage of overlap in time between Target and Distractor VM calls.

#### Pulse-Number Index

Custom software extracted all phee calls produced by subjects in each test session and identified the number of pulses within these calls based on previously identified stereotyped spectro-temporal structure of these vocalizations ^32^. Once cataloged, we then compared the number phee calls produced that comprised 1, 2 or 3+ pulses. Previous studies have shown that the modal marmoset phee variant consist of 2-pulses, while the other variants occur at lower frequency ^32^. Phee calls consisting of 3 or more pulse calls were rarely produced in the current experiments, accounting for <10% of calls, these were grouped together. Because the number of phee calls comprising 3+ pulses did not vary across the test conditions, these were excluded from this this metric. We generated the Pulse-Number Index by calculating the difference over the sum of the 1 and 2 pulsed phee calls produced in each session [(1*PulseRatio* – 2*PulseRatio*)/(1*PulseRatio* + *2PulseRatio*)]. Positive values would indicated a bias towards 1-Pulse Phee calls, while a negative value reflects a bias towards 2-pulse Phee Calls.

#### Estimated PDF

The estimated PDF was calculated using the MATLAB function ksdensity with Kernel set to normal, function to PDF, Boundary Correction to reflection, and the support set at 0 to the maximum value found in the distribution for a given plot. Confidence intervals within Estimated PDFs (Figure 2A,C) were created by getting a ksdensity plot at the same support boundaries for each session for a given distribution and then finding the mean and 95% confidence intervals for the same x-positions.

#### Cumulative counts

For Figure 2B,D, we subdivided each recorded session into one second bins and counted how many events occurred for the required analysis in each bin. Then we took the cumulative sum and divided it by the sum for each session to get the normalized plots. Each session’s cumulative distribution for the respective data was then put collapsed by mean and 95% confidence intervals. Preliminary tests showed a normal distribution for each respective bin.

### Latency in conversation

As mentioned previously, conversations were defined by two or more consecutive responses by the subject to the target within the antiphonal delay (10 sec). All calls produced by the subject and Target VM that occurred within call exchanges that had at least 2 subject responses was included. In cases where the subject initiated the conversation, a third subject call would be needed to be included. The latency of the subject to respond within the sequence of exchanges was used for analysis.

### Linear Model Analysis

The following were input into the Linear Model – VM Pulse # (2-pulse:Expt 1, 1-pulse:Expt2), Low and High Distractor Density, and Fixed and Single conditions – for a total of 144 sessions. We also chose to include the calculated Distractor Density for each session along with the Distractor ICI (see Methods for more details on setup). Given a strong positive correlation between Distractor ICI and standard deviation (Pearson’s Linear Correlation: rho = 0.931 and p < 0.0001), we took the coefficient of variance (COV, standard deviation divided by mean) as a way to encapsulate these two correlated factors while avoiding rank deficiency in any linear model (COV v Mean ICI, Pearson’s Linear Correlation rho = −0.0956, p = 0.254. Figure 5B). This also gave an added benefit of enumerating the relative dispersion of the Distractor ICI. This analysis yielded six total predictor variables. The following 8 vocal behavior response variables were also inputted into the model: the mean duration of all calls, the duration of the 1-Pulse calls, Index of relative 1 and 2 pulse calls produced by subjects (Pulse Number Index), the mean Communication Index, subjects mean latency to respond in a conversation, the number of subject calls produced, the number of conversations, and the mean length of those conversations. We performed analyses only on the Fixed-Location and Single-Location conditions because the Random-Location was not performed in Experiment 2.

**Figure 5.**
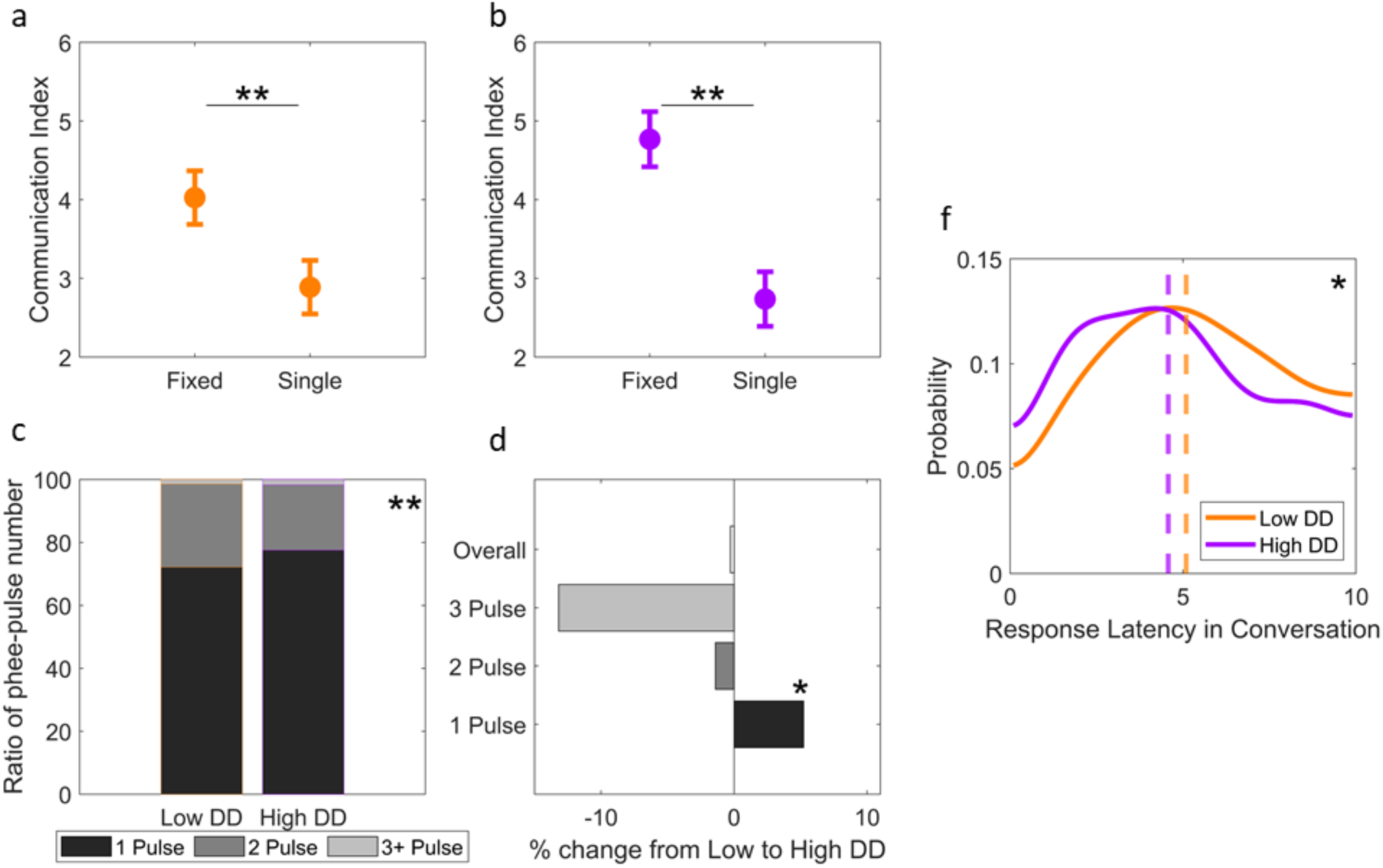
Results from Experiment 2. (a, b) Plots the Mean Communication Index [95% CI] for Fixed-Location and Single-Location test conditions. ** Significant difference between two conditions, p < 0.0001. (a) Plots Communication Index for the Low Distractor Density condition in red, while (b) plots the High Distractor Density condition in blue. (c) Stacked bar graph showing the distribution phee calls produced by subjects that comprised 1-Pulse (black), 2-Pulses (dark grey) and 3 or more pulses (light grey) in both the Low and High Distractor Density (DD) environments. ** Significant difference between distributions, p < 0.0001 (d) The change in duration of the phee calls comprising 1, 2, 3 and Overall duration is shown as percent change from Low DD to High DD conditions. * Significant difference for that category, p < 0.001 (e) Probability density estimate plots of subjects’ latency to respond to the Target VM in conversations in both Low DD (red) and High DD (blue) conditions. The median value is shown as a dashed, vertical red line – Low DD – and blue line – High DD. * Significant difference for that category, p < 0.001.

**Figure 6.**
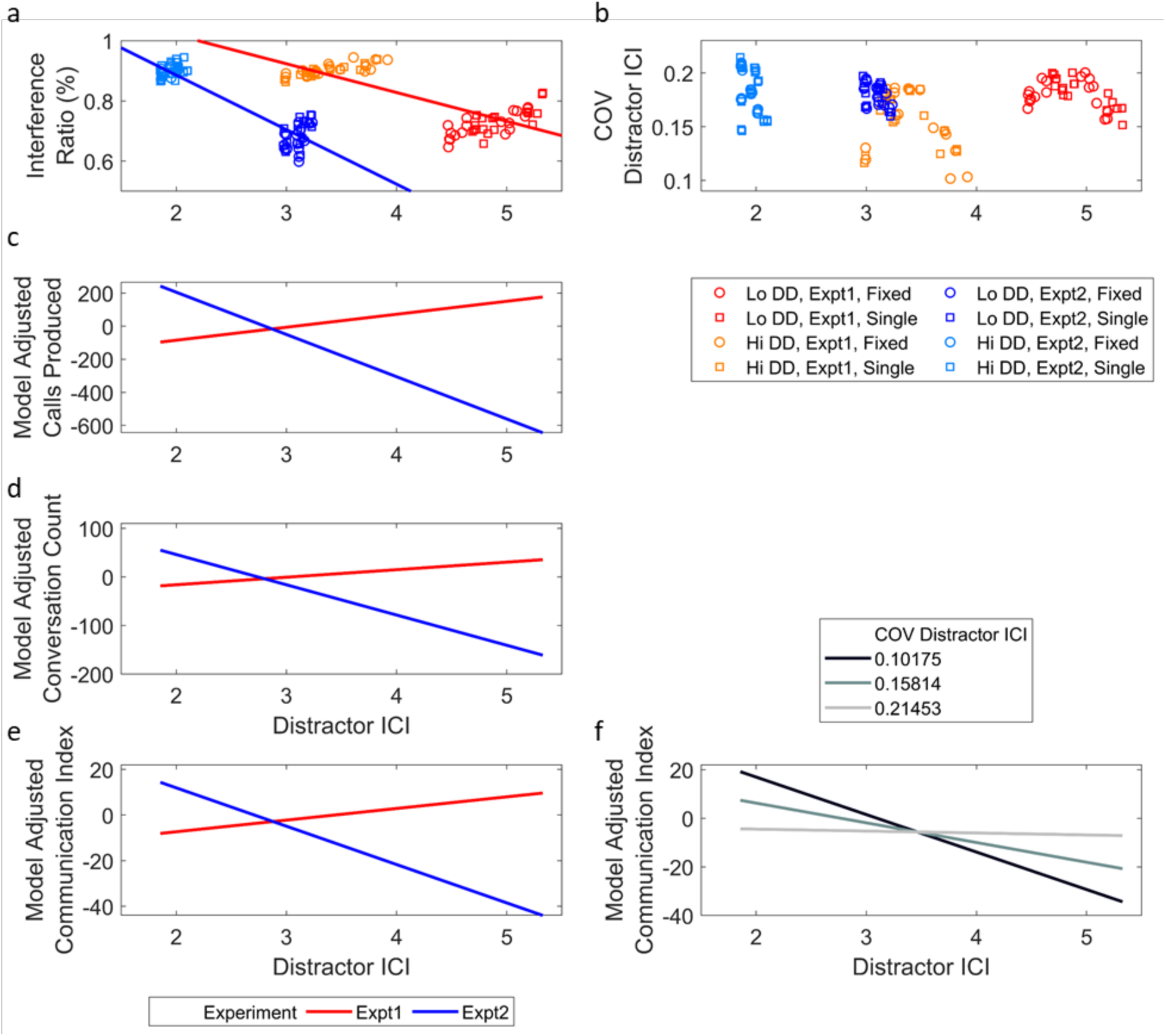
Linear Model Outcome (a) Scatter plot displaying Interference Ratio for the Distractor ICI measured during in each test session. Lines represent the least-squares fit for each Experiment. (b) Plots the COV Distractor ICI for the Distractor ICI measured during in each test session. Figure legend for (a & b) is shown below (b). (c-e) Significant interactive effects of Distractor ICI with different metrics of vocal behavior revealed by the linear model are shown. Results of the model from Experiment 1: 2-pulse VM phee calls (red line) and Experiment 2: 1-pulse VM phee calls are shown (blue line). The adjusted response value accounts for the average values of all other terms except Distractor ICI x Experiment within the linear model. (c) Plots Distractor ICI cross Experiment type by the adjusted response variable of Calls Produced by the subject in each session. (d) Plots Distractor ICI cross Experiment type by the adjusted response variable of Conversation Count. Conversation Count refers to number of conversations with at least two or more subject responses made in a session. (e) Plots the relationship between model adjusted median Communication Index of the subject for Target VM by the Distractor ICI. (f) Distractor ICI x COV Distractor ICI term (which represents the standard deviation of Distractor ICI) is plotted against its effect on the Conversation Index. COV values plotted include minimum (light grey), maximum (dark grey), and the average of the two (mid-grey).

#### Response Variables

These metrics were used as response variables within our linear models as mentioned in the Results section. Each one was calculated for each recorded session within a given experimental condition (18 per condition).

- *Average Duration of Calls:* The mean duration of subject calls.
- *Duration of 1 Pulse Calls:* Mean duration of 1-pulse calls produced by the subject
- *Pulse-Number Index:* The difference over sum of the ratio of one pulse calls to two pulse calls produced by the subject.
- *Communication Index:* The mean position of the subject calls as previously mentioned.
- *Response Latency in Conversation:* The mean latency of subjects to respond to Target VM within a conversational exchange.
- *Number of Calls:* Number of calls produced by the subject in a given session.
- *Number of Conversations:* The number of times the subject engaged in conversational exchanges.
- *Length of Conversations:* The mean number of subject calls produced within each conversation.

#### Design

MATLAB function ‘fitlm’ was used to fit six predictor variables to each of the 8 response variables thus creating 8 linear models of comparison on 144 observations per model. The six predictor variables were: the calculated Distractor Density (as seen in Figure 3B,C and Figure 4A,B), Distractor ICI, COV Distractor ICI, the categorical Distractor Density (Low or High), the categorical spatial configuration (Fixed or Single), and the categorical Experiment (2-Pulse or 1-Pulse). An interactive linear model was created that included an intercept term (1), linear term for each predictor (6), and products of pairs of distinct predictors excluding squared terms (15), for a total of 22 predictor terms. The 8 models created with 22 predictor terms were corrected for multiple comparisons using the Bonferroni correction. With a criterion at α = 0.05, the new p-value threshold was calculated to be at 0.05/176 = 0.000284. Any model’s F-test for a degenerate constant model that was below this threshold was included for further analysis of the terms. Three models reached this threshold as mentioned in the results. Of those three, only terms with coefficients that were significantly different from 0 below the corrected new significance threshold were subsequently explored in Figure 5C-G.

#### Test of Assumptions

The significant models’ residuals were finally looked at to test for homoscedasticity and normality of the residuals. All three initial models (Number of Calls, Number of Conversations, Communication Index) failed the normal distribution (p=0.0125, 0.0179, 0.001), but the homoscedasticity was preserved in the models (Breusch-Pagan test, df=6, p =.4724, 0.0603, 0.6832). Looking at the normal plots, there was clear evidence of some outliers in the data. Taking the residuals from the Communication Index model, we removed residual outliers 1.5 times outside the quartiles at 25% and 75% of the data. Of the 144 points, eight points were outliers along with 3 NaNs (7.64%) that were removed. After removal, the same three models were once more analyzed. The reported final values in the Results section indicate these new values. All three had normal distributions of the residuals as indicated by a failure to reject the null hypothesis of normality by the Lilliefors test (p=0.270, 0.111, 0.0684). As well, the model for Communication Index and Number of Calls failed to reject the null-hypothesis of homoscedasticity in the Bresuch-Pagan test with studentized Koenker’s statistic (Breusch-Pagan test, df=6, p=0.147,0.959), while the model for Number of Conversations was on the threshold (p = 0.0406). We finally looked at the collinearity of the predictor variables and found that of the three continuous variables (Distractor ICI, COV Distractor ICI, and Distractor Density), none of them exhibited multicollinearity as determined by the Belsley collinearity diagnostics (Condition indeces for the three 1, 5.96, 12.3).

## Data Availability

The data generated during the experiment along with the associated analyses and figure creations done for the paper have been deposited in Dryad with the primary access to create the figures and the statistical tests mentioned in the manuscript can be found in the Dryad repository with the identifier doi:10.6076/D1RG6V and can be permanently found at this link https://datadryad.org/stash/dataset/doi:10.6076/D1RG6V

## Acknowledgements

We thank Victoria Ngo and Madeline Gagne for assistance in data collection and Drs. Yi Zhou and Vatsun Sadagopan for comments and discussion on an earlier draft of this manuscript. This work supported by grants from NIH (R01 DC012087) and DARPA (SSC-5029) to CTM.

## Ethics Statement

This study was performed in strict accordance with the recommendations in the Guide for the Care and Use of Laboratory Animals of the National Institutes of Health. All of the animals were handled according to approved institutional animal care and use committee (IACUC) protocols and approved by the University of California San Diego (#S09147).

## Author Contributions

VJ designed the experiments, collected the data, analyzed the data and wrote the manuscript. CM designed the experiments, oversaw data collection and analysis and wrote the manuscript.

The Authors declare no competing interests.

